# Cellular calcium levels influenced by NCA-2 impact circadian period determination in *Neurospora*

**DOI:** 10.1101/2021.05.24.445544

**Authors:** Bin Wang, Xiaoying Zhou, Scott A. Gerber, Jennifer J. Loros, Jay C. Dunlap

**Author notes:** Address correspondence to Jay C. Dunlap,. Bin Wang and Xiaoying Zhou contributed equally to this work and Bin Wang completed the project and wrote up the manuscript so listed first.

## Abstract

Intracellular calcium signaling has been implicated in control of a variety of circadian processes in animals and plants but its role in microbial clocks has remained largely cryptic. To examine the role of intracellular Ca^2+^ in the *Neurospora* clock we screened knockouts of calcium transporter genes and identified a gene encoding a calcium exporter, *nca-2*, uniquely as having significant period effects. Loss of NCA-2 results in an increase in cytosolic calcium level, and this leads to hyper-phosphorylation of core clock components, FRQ and WC-1, and a short period as measured by both the core oscillator and overt clock. Genetic analyses showed that mutations in certain *frq* phospho-sites, and in *Ca*^*2+*^*-calmodulin-dependent kinase* (*camk-2*), are epistatic to *nca-2* in controlling the pace of the oscillator. These data are consistent with a model in which elevated intracellular Ca^+2^ leads to increased activity of CAMK-2 leading to enhanced FRQ phosphorylation, accelerated closure of the circadian feedback loop, and a shortened circadian period length. At a mechanistic level some CAMKs undergo more auto-phosphorylations in Δ*nca-2*, consistent with high calcium in the Δ*nca-2* mutant influencing the enzymatic activity of CAMKs. NCA-2 interacts with multiple proteins including CSP-6, a protein known to be required for circadian output. Most importantly, expression of *nca-*2 is circadian clock-controlled at both the transcriptional and translational levels, and this in combination with the period effects seen in strains lacking NCA-2, firmly places calcium signaling within the larger circadian system where it acts as both an input to and output from the core clock.

**Importance:** Circadian rhythms are based on cell-autonomous, auto-regulatory, feedback loops formed by interlocked positive and negative arms, and they regulate myriad molecular and cellular processes in most eukaryotes including fungi. Intracellular calcium signaling is also a process that impacts a broad range of biological events in most eukaryotes. Clues have suggested that calcium signaling can influence circadian oscillators through multiple pathways; however, mechanistic details have been lacking in microorganisms. Building on prior work describing calcium transporters in the fungus *Neurospora*, one such transporter, NCA-2, was identified as a regulator of circadian period length. Increased intracellular calcium levels caused by loss of NCA-2 results in over-activation of calcium-responsive protein kinases, in turn leading to a shortened circadian period length. Importantly, expression of NCA-2 is itself controlled by the molecular clock. In this way calcium signaling can be seen as providing both input to and output from the circadian system.

## Introduction

In most eukaryotes and certain prokaryotes, circadian clocks link environmental cues such as temperature and light to metabolism to regulate various physiological and molecular events ranging from virulence and immunity to cell cycle control (1–3). In fungi and mammals, the core circadian machinery is built based on a transcriptional-translational feedback mechanism in which the positive arm drives transcription of components comprising the negative arm which, in turn, feeds back to repress the positive arm, terminating its own expression. *Neurospora crassa* has been widely used as a model eukaryote for circadian studies for decades. In *Neurospora*, the White Collar Complex (WCC), formed from WC-1 and WC-2, serves as the positive arm transcriptional activator for the core clock gene *frequency* (*frq*) by binding to one of two DNA elements, the *Clock box* (*C box*) in the dark or the *Proximal Light-Response Element* (*PLRE*) in the light (4–6). FRQ, the gene product of *frq*, interacts with FRH (FRQ-interacting RNA Helicase) (7, 8) and CK1 (casein kinase 1) (9) to form the FFC complex, the negative arm that represses WCC activity by promoting its phosphorylation at a group of residues (10).

Protein phosphorylation has been shown to control protein functions via protein-protein/DNA associations, protein stability and activity, and subcellular localization, all of which have been proven or suggested to regulate functions of circadian components (11–14). In *Neurospora*, FRQ is intricately regulated by over 100 time-specific phosphorylation events (9, 15); multiple kinases, such as CKI, CKII, PKA, and CAMK-1, and phosphatases, like PP2A, have been reported to directly or indirectly control FRQ phosphorylation status (16–18). Extensive phosphorylation has also been observed on WCC in light and dark conditions (10, 16, 19, 20). Recently, over 90 phosphoresidues have been mapped on WC-1 and WC-2, governing their circadian repression and controlling circadian output, and a small subset of these has been shown to be essential for the feedback loop closure (10).

Calcium as a second messenger regulates a wide variety of cellular pathways: for example, elevated Ca^2+^ in the cytosol and mitochondria of neurons is required to synchronize neuronal electrical activity (e.g., reviewed in (21)); all muscle fibers use Ca^2+^ as their main regulatory and signaling molecule (e.g., reviewed in (22)); during mammalian fertilization, Ca^2+^ influx induces oocyte development in many species (23). At the molecular level, enzymes and other proteins can be regulated by calcium via allosteric regulatory effects (24). Indeed, diverse evidence also connects calcium signaling with circadian regulation. In *Arabidopsis thaliana*, the concentration of cytosolic Ca^2+^ oscillates over time (25, 26), which regulates circadian period length through action of a CALMODULIN-LIKE protein on the core circadian oscillator (27). Circadian oscillation of Ca^2+^ has been observed in hypothalamic suprachiasmatic nucleus (SCN) neurons, driving daily physiological events (28). In addition a small body of literature has described effects of calcium ionophores and calmodulin antagonists on the *Neurospora* clock (29–33). Although this research was published before there was sufficient understanding of basic cellular physiology to fully interpret the work, it provides a rich context for studies on the role of calcium signaling in the *Neurospora* clock.

Despite the paucity of recent data on circadian effects of calcium in fungi, the cellular physiology of calcium metabolism in fungi including *Neurospora* is well understood (34–40) and is consistent with general knowledge of animal cells. The resting concentration of Ca^2+^ in the cytoplasm of fungal and mammalian cells is normally maintained at 50-200 nM (41–45), which is 20,000- to 100,000-fold lower than that in a typical extracellular environment (46). To be maintained at this low level in the cell, Ca^2+^ is actively pumped out from the cytosol to the extracellular space, reticulum, vacuole, and/or mitochondria (34, 35, 47–51); bearing binding affinity to Ca^2+^, certain proteins in the cell can also contribute to lowering the level of free cytosolic Ca^2+^ (52).

To elicit signaling events, the cell releases Ca^2+^ from organelles or Ca^2+^ enters the cell from extracellular environments. When stimulated by certain signals, cytoplasmic Ca^2+^ can be suddenly increased to reach ∼500-1,000 nM through activation of certain ion channels in the ER and plasma membrane or indirect signal transduction pathways, such as G protein-coupled receptors (e.g., reviewed in (53, 54)). Cytosolic calcium bursts lead to activation of Ca^2+^–calmodulin-dependent protein kinases (CAMKs) (55–59). In mammals, the CAMK cascade includes three kinases: CaM-kinase kinase (CaMKK), CaMKI, and CaMKIV; CaMKI and CaMKIV are phosphorylated and activated by CaMKK (55, 60–65). CaMKK and CaMKIV reside in the nucleus and cytoplasm while CaMKI is only located in cytosol; nuclear CaMKIV promotes phosphorylation of several transcription factors, such as CREB and CBP, to regulate gene expression (60, 66, 67). The *Neurospora* genome encodes four CAMK genes that are subject to diverse regulation, although little is known about their intracellular localization (18, 37).

By impacting a wide range of cellular processes, circadian clocks and calcium signaling are two classic regulatory mechanisms evolved to coordinate environmental factors, cellular responses, and metabolism. In this study, a screen of calcium regulators identified *nca-2*, a calcium pump gene, as a regulator of circadian period length in *Neurospora*. In Δ*nca-2* strains, FRQ and WC-1 become hyper-phosphorylated; deletion of *camk-2* individually blocks the period shortening effect and FRQ hyper-phosphorylations in Δ*nca-2*. NCA-2 interacts with multiple proteins, which suggests that it might function in other cellular processes in addition to the circadian clock.

## Results

### Identification of *nca-2* as a regulator of the *Neurospora* circadian clock

Calcium signaling impacts circadian processes (e.g., 18, 30, 31) and directly controls a wide range of cellular and physiological events, but the means through which it impacts the circadian system is not fully described. *Neurospora* encodes several calcium transporter genes including *nca-1* (a SERCA-type ATPase), two closely related genes, *nca-2* and *nca-3* (PMCA-type ATPases), *pmr-1* (a SPCA-type Ca^2+^-ATPase), and *cax* (a vacuolar Ca^2+^/H^+^ exchanger (**35**)). To facilitate monitoring circadian phenotypes, individual knockouts of these calcium pump genes were backcrossed to *ras-1*^*bd*^ and *frq C box*-driven *luciferase* and analyzed by race tube and luciferase assays. Of these deletion mutants tested, Δ*pmr-1* shows an extremely slow growth rate on race tubes (Figure 1A) but is nicely rhythmic with a slightly shorter period in the luciferase assay (Figure 1B); disruption of *nca-2*, a plasma membrane-located calcium pump, leads to a ∼2-hr shorter period than that in wild type (WT) by race tube (Figure 1A) and luciferase (Figure 1B) analyses. (Of note, although on any given day the period estimates of strains bearing mutated calcium pumps showed normal precision, period length assays done on different days were more variable than is typical. For this reason, comparisons within figures always reflect assays of different strains done on the same day with the same medium.) Appearing at DD12, newly synthesized FRQ in Δ*nca-2* is slightly more abundant than that in WT (Figure 1C, left) and *frq* mRNA levels in Δ*nca-2* are substantially higher in the subjective circadian night phase (DD4, 8, 24, 28) as compared to WT (Figure 1C, right), consistent with a faster running circadian clock in Δ*nca-2* (Figure 1A and 1B). The cytosolic calcium level in Δ*nca-2* is increased about 9.3 fold as compared to WT (36) suggesting a basis for this period change. To verify that the period shortening in Δ*nca-2* was due to this increased intracellular Ca^2+^, the Δ*nca-2* strain was examined on race tubes prepared without calcium in the medium. Interestingly, Δ*nca-2* displays a WT period on race tubes using Ca^2+^-free medium while its clock becomes ∼4-hr shorter than WT with normal levels of calcium in the medium (Figure 1D), confirming that the role of *nca-2* in regulating the pace of the circadian oscillator is through controlling the cytosolic calcium level. These data indicate that *nca-2* is required for keeping calcium in cytosol at reduced levels to maintain a normal circadian period.

**Figure 1.**
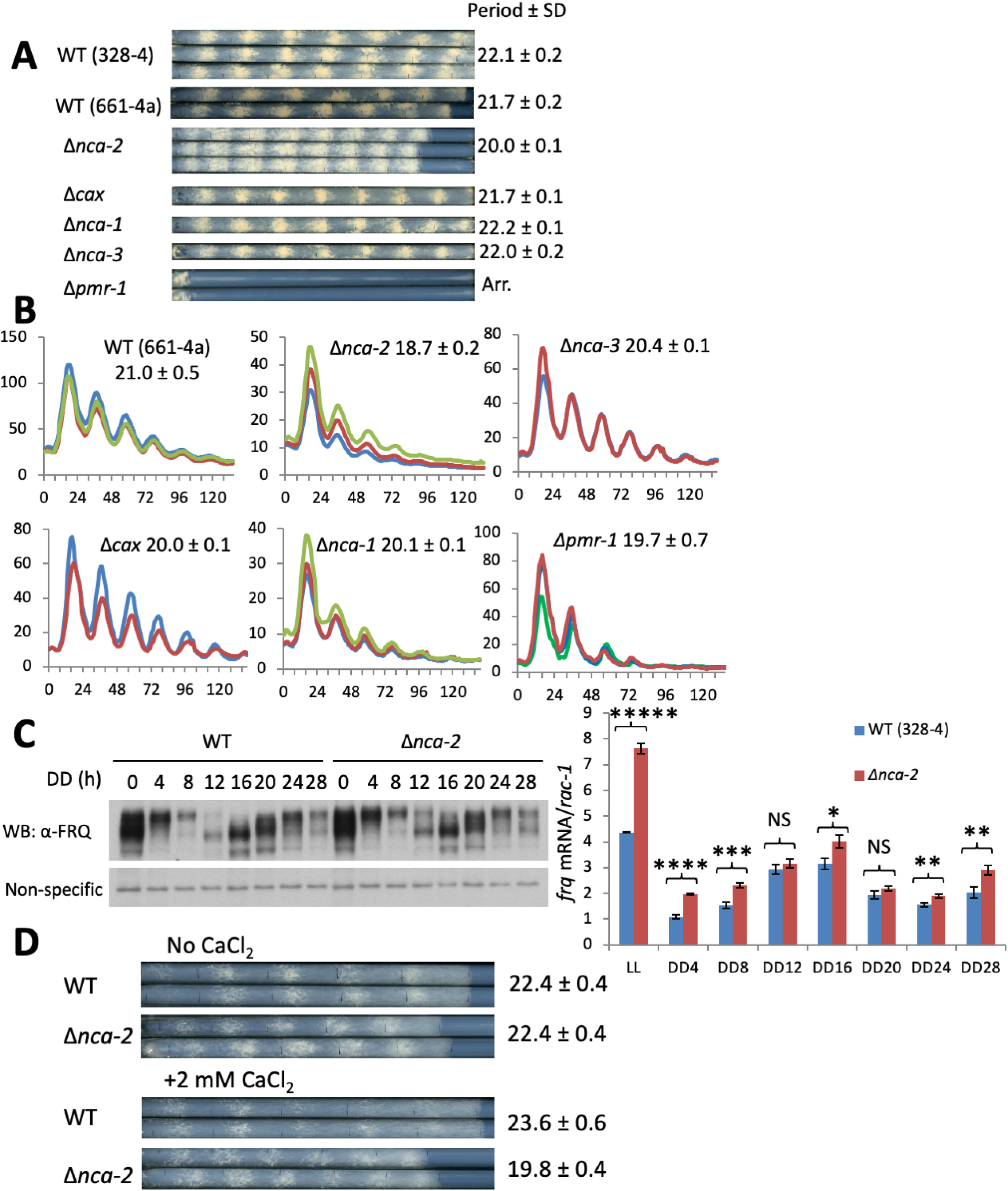
Gene deletions of calcium pumps were tested for circadian phenotypes by race tube (**A**) and luciferase (**B**) analyses. Strains were cultured on 0.1% glucose race tube medium in a 96 well plate and synchronized by growth in constant light overnight (16-24 hrs) followed by transfer to darkness. Bioluminescent signals were monitored by a CCD-camera every hour, bioluminescence data were acquired using ImageJ with a custom macro, and period lengths were manually calculated. Raw bioluminescent data from three replicates were plotted with the X-axis and Y-axis representing time (hrs) and arbitrary units respectively. (**C**) Left, Western blot showing the expression level of FRQ in WT and Δ*nca-2* over 28 hr detected with FRQ-specific antibody. DD, hours after the light to dark transfer; right, RT-qPCR showing relative levels of *frq* mRNA expressed in WT and Δ*nca-2*; *rac-1* was used as an internal control, to which *frq* expression is normalized (n = 3, mean values ± standard error of the mean). Asterisks indicate statistical significance when compared with WT as determined by a two-tailed student’s *t* test. *****P<0.00001; ****P=0.00006; ***P=0.001337; **P<0.01; *P=0.010131. NS, difference is not significant. (**D**) Race tube assays of WT and the Δ*nca-2* strain using race tube media in the presence or absence of 2 mM calcium chloride. Growth fronts of the strains were marked by vertical blacklines every 24 hr. *nca-3* (NCU05154): calcium P-type ATPase; *nca-1* (NCU03305): calcium-transporting ATPase sarcoplasmic/endoplasmic reticulum type; *cax* (NCU07075): calcium/proton exchanger; *pmr-1* (NCU03292): calcium-transporting ATPase type 2C member 1; *nca-2* (NCU04736): plasma membrane calcium-transporting ATPase 3; gene names, NCU numbers, and descriptions were obtained from the FungiDB website (https://fungidb.org/fungidb/app). Period was determined as described in Material and Methods and is reported ± SD (n=3).

### WC-1 and FRQ are hyper-phosphorylated in Δ*nca-2*

WC-1 and FRQ are essential components in the positive and negative arms respectively of the *Neurospora* feedback loop and their phosphorylation has been proven to play an essential role in the determining their circadian functions (9, 10, 15, 16, 19). In addition to serving as the main transcription factor driving expression of *frq*, WC-1 is also the principal blue light photoreceptor for the organism, forming a homodimer (4) and getting hyper-phosphorylated (20) upon light exposure. To probe WC-1 and FRQ in Δ*nca-2*, amounts and phosphorylation profiles of WC-1 and FRQ were analyzed by Western blotting using specific antibodies. The stability of FRQ in Δ*nca-2* is very similar to that in WT (Figure S1), and although WC-1 appeared slightly less stable, the cellular levels of WC-1 were even above those of WT, altogether suggesting that the stability of the core clock components does not determine the shortened period in Δ*nca-2* and the WC-1 level and stability are not consistent with the period length shortening in Δ*nca-2*. Following a light-pulse, WC-1 is more abundant and hyperphosphorylated in Δ*nca-2* as compared to WT (Figure 2A) whereas, surprisingly, expression of *wc-1* is significantly lower than that in WT (Figure 2B); consistent with the data from the light-pulse experiment, in the dark, Δ*nca-2* contains a higher level of WC-1 with more phosphorylations (Figure 2C) despite a low mRNA level (∼20-50% as in WT) (Figure 2D). These data suggest that *nca-2* regulates *wc-1* expression at both transcriptional and post-transcriptional levels independent of light and dark conditions. The hyper-phosphorylation of WC-1 in Δ*nca-2* was confirmed by a more sensitive assay (Figure 2E) using Phos-tag gels (68) such as have been applied to resolve single phosphoresidues on WC-1 and WC-2 (10). Similar to WC-1, FRQ in Δ*nca-2* is also more heavily phosphorylated than that in WT at DD14, 16, and 18 (Figure 2F) when newly synthesized FRQ is the dominant form in the cell, and at DD24 (Figure 2G) when all FRQ becomes extensively phosphorylated prior to its turnover (see also Figure 1A). Altogether, these data demonstrate that WC-1 and FRQ become hyper-phosphorylated in Δ*nca-2*, suggesting that the elevated calcium in Δ*nca-2* might lead to over-activation of kinase(s) or repression of phosphatase(s) targeting FRQ and WC-1, thereby altering their activities in the clock.

**Figure 2.**
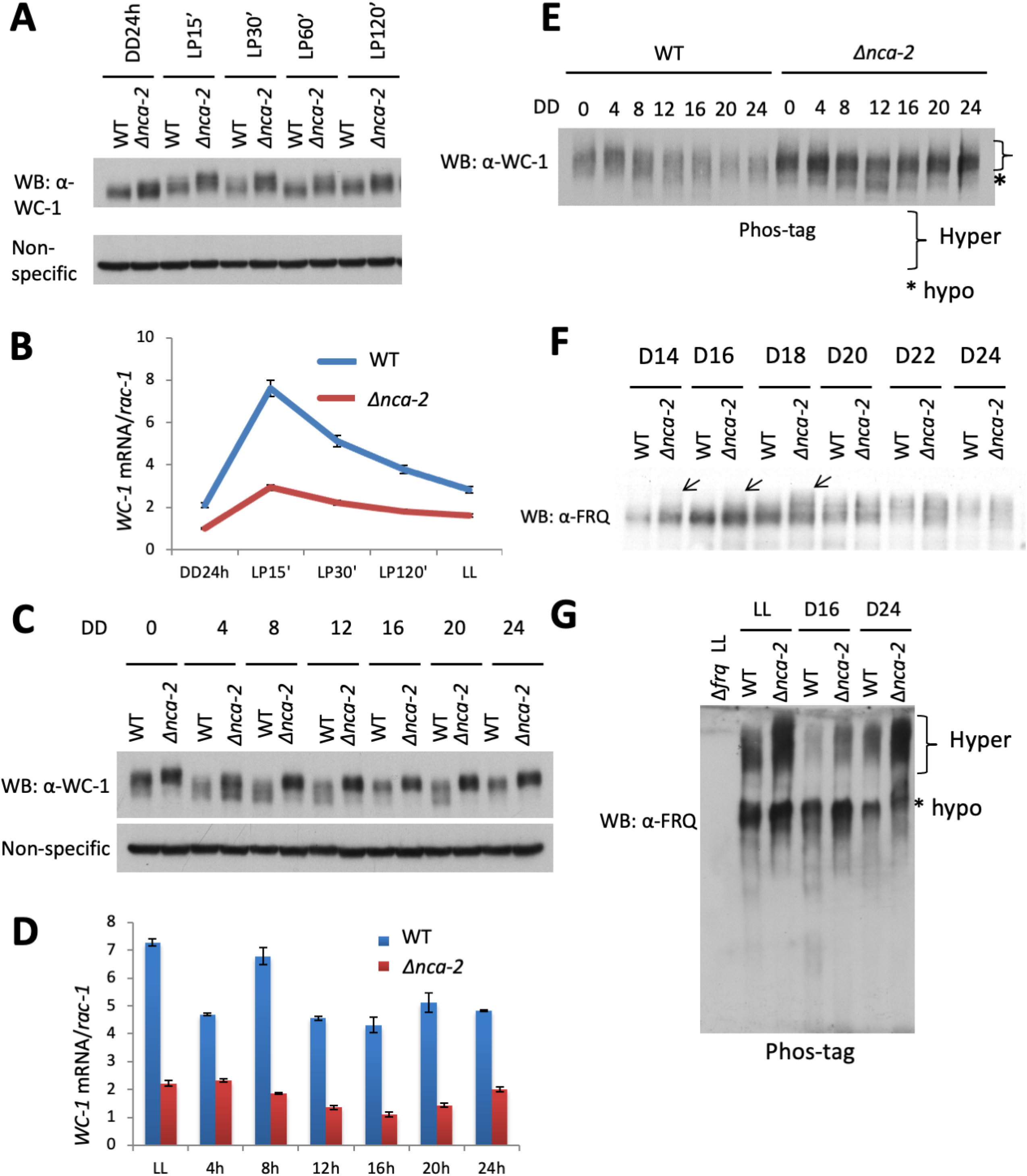
Circadian components WC-1 and FRQ are hyper-phosphorylated in Δ*nca-2*. (**A**) Total WC-1 was followed by Western blotting. Samples were cultured in the constant dark prior to a 15, 30, 60, and 120 min light exposure. Non-specific bands in the same blot were shown for equal loading. Decreased electrophoretic mobility is indicative of phosphorylation status (7). (**B**) mRNAs extracted from samples cultured in the dark for 24 hrs or following a 15, 30, or 120 min light exposure as indicated were reverse-transcribed to cDNA followed by quantitative PCR with a primer set specific to *wc-1*. (**C**) Western blotting of WC-1 in a 24-hr time course with a 4-hr interval. (**D**) Similar to (**C**), RT-qPCR was performed with samples harvested at the circadian conditions indicated. Phosphorylation profiles of WC-1 (**E**) and FRQ (**F, G**) in WT and Δ*nca-2* were analyzed by Western blotting using SDS-PAGE gels bearing (G) 20 μM Phos-tag chemicals and a ratio of 149:1 acrylamide to bis-acrylamide. (**F**) Western blotting of FRQ in WT and Δ*nca-2* from DD14 to DD24 with a 2-hr resolution. * in **E** denotes the mobility of unphosphorylated WC-1, and the bracket the region corresponding to hyperphosphorylated WC-1. Arrows indicate hyper-phosphorylated FRQs observed in Δ*nca-2*.

### Epistasis analysis is consistent with an effect of Δ*nca-2* on FRQ but not on WCC

FRQ is phosphorylated in a time-specific manner at over 100 sites, and elimination of certain phospho-sites in different domains can cause opposite phenotypes on period lengths (9, 15). Because loss of *nca-2* elicits FRQ hyperphosphorylation at almost all time points examined (Figure 2F and 2G), we reasoned that this enhanced FRQ phosphorylation in Δ*nca-2* might contribute to the short period length in this strain. If this is so, then circadian period lengths in *frq* mutants encoding proteins that cannot be phosphorylated at key residues should not display the period shortening. To this end, several *frq* phospho-mutants displaying long circadian periods from (9) were individually backcrossed to Δ*nca-2* and *frq-luc*, and assayed by tracking bioluminescent signals in real-time in darkness. While circadian periods of *frq*^*S541A, S545A*^ and *frq*^*S548A*^, and *frq*^*7*^ responded to loss of *nca-2* as did WT (Figure 3 and Figure S2A), absence of *nca-*2 does not significantly influence the period length of *frq*^S72A, *S73A, S76A*^, *frq*^*S538A, S540A*^, or *frq*^*S632A, S634A*^ (Figure 3). These are all residues whose inability to be phosphorylated results in period lengthening (9), so the epistasis of these *frq* alleles is consistent with NCA-2 influencing FRQ phosphorylation at these sites.

**Figure 3.**
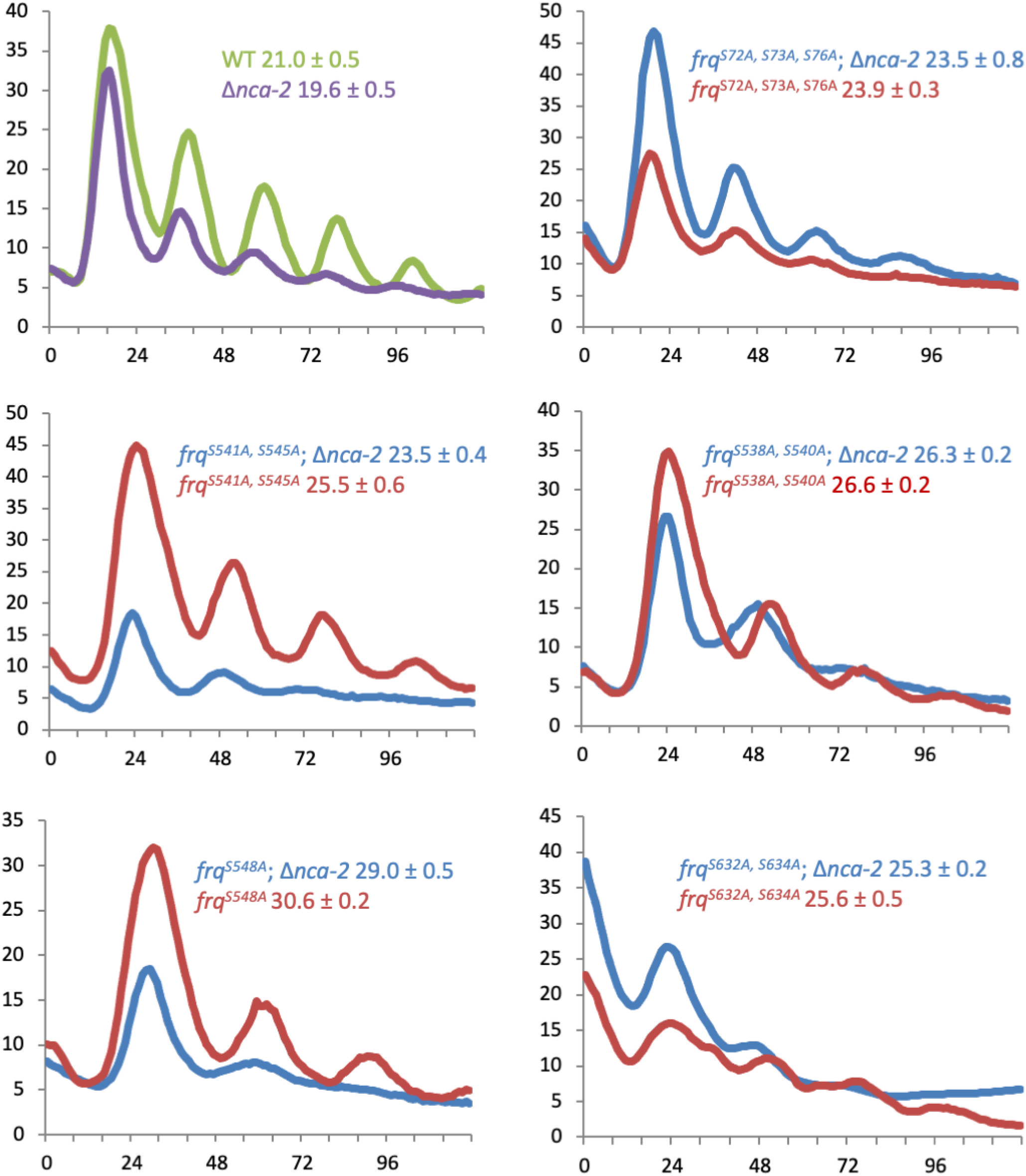
Some *frq* alleles are epistatic to Δ*nca-2*. The *frq C box* promoter activity was measured using *C box-luciferase* at the *his-3* locus in indicated *frq* phospho-mutants in the presence or absence of *nca-2*. Strains were grown on 0.1% glucose race tube medium in constant light overnight (16-24 hrs) prior to transfer to darkness. *frq*^*S72A, S73A, S76A*^, *frq*^*S541A, S545A*^, *frq*^*S538A, S540A*^, *frq*^*S548A*^, and *frq*^*S632A*,^ S634A were derived from (9). Period was determined as described in Material and Methods and is reported ± SD (n=3).

To examine the effect of Δ*nca-2* on WCC phosphorylation and period length in the same manner, Δ*nca-2* was backcrossed to several *wcc* mutants eliminating key phosphoresidues that have been identified and shown to determine the circadian feedback loop closure (10), and monitored by the luciferase assay. The absence of *nca-2* further shortens the periods of *wc-1*^*S971A, S988A, S990A, S992A, S994A*,^ *S995A* and *wc-2*^*S433A*^ respectively (Figure S2A), suggesting that *nca-2* regulates the core oscillator independent of WCC phosphorylation at the sites essential for its repression. Consistent with this, in Δ*nca*-2 the phosphorylation levels of WC-1 S971 and S990, two key sites required for FFC-mediated WCC repression, are similar to that in WT (Figure S2B), further suggesting that altered phosphorylation of the positive arm in the oscillator is not the cause of the short period of Δ*nca-2*.

### Δ*camk-2* does not further shorten the period of Δ*nca-2*

Data in Figures 2 and 3 are consistent with NCA-2 acting through kinases or phosphatases on FRQ, and the elevated calcium in Δ*nca-2* (36) might activate Ca^2+^-responsive kinases to over-phosphorylate FRQ (Figure 2F and 2G). CAMKs have been well documented to be activated by the elevated intracellular Ca^2+^ and calmodulin. There are four *camk* genes (*camks-1-4*) annotated in the *Neurospora* genome and their catalytic domains are conserved despite a low overall identity of amino acid sequences (37). Expression of *camk-1-4* genes moderately increases in Δ*nca-2* compared to those in WT across 28 hrs in the dark (Figure S3). Among the four CAMKs, CAMK-1 has been reported to directly phosphorylate FRQ at multiple sites *in vitro* although only a very subtle period defect was observed in Δ*camk-1* (18); however, in our hands the Δ*camk-1* strain showed greatly reduced growth and was arrhythmic on race tubes (Figure S4A), suggesting that prior data may have reflected a revertant strain. To further evaluate this and characterize roles for CAMKs we made all combinations of Δ*camk* mutants, transformed these with the *C-box luc* reporter, and assayed their clocks. We found that circadian periods in individual or combinational knockouts of *camks* are indeed quite similar to WT (Figure S4B). To test whether Δ*nca-2* regulates the clock through *camks-1-4*, Δ*nca-2* was backcrossed to Δ*camks-1-4* respectively and circadian periods were assayed by luciferase analyses. Interestingly, Δ*camks-1, -3, and -4* each showed the characteristic period shortening when in combination with Δ*nca-2*; however, Δ*camk-2* Δ*nca-2* showed the same circadian period as the Δ*camk-2* single mutant with no additional shortening due to Δ*nca-2* (Figure 4A), suggesting that *nca-2* and *camk-2* function in the same pathway to regulate the circadian period. Because in certain cases activated kinases not only phosphorylate their substrates but also actuate autophosphorylation in *cis* or in *trans*, phosphorylation on these kinases can be indicative of their activities. To test this, the phosphorylation status of CAMKs-1-4 were measured by Western blotting using the 149:1 (acrylamide/bisacrylamide) Phos-tag gel that has been used to resolve single phosphorylation events on WC-1 and WC-2 (10). CAMKs-2 and -4 display similar phospho-profiles in the presence or absence of *nca-2*, while, interestingly, CAMKs-1 or -3 in Δ*nca-2* undergo more phosphorylations than those in the WT background (Figure 4B), suggesting that their activities might be stimulated due to elevated calcium resulted from the absence of *nca-2*. Taken together, these data suggest that the elevated calcium concentration in Δ*nca-2* directly or indirectly activates CAMKs, which leads to hyper-phosphorylation of FRQ, thereby shortening the circadian period. The data further indicate that although intracellular calcium can influence periodicity through CAMKs, phosphorylation by CAMKs is not required for rhythmicity; it is modulatory.

**Figure 4.**
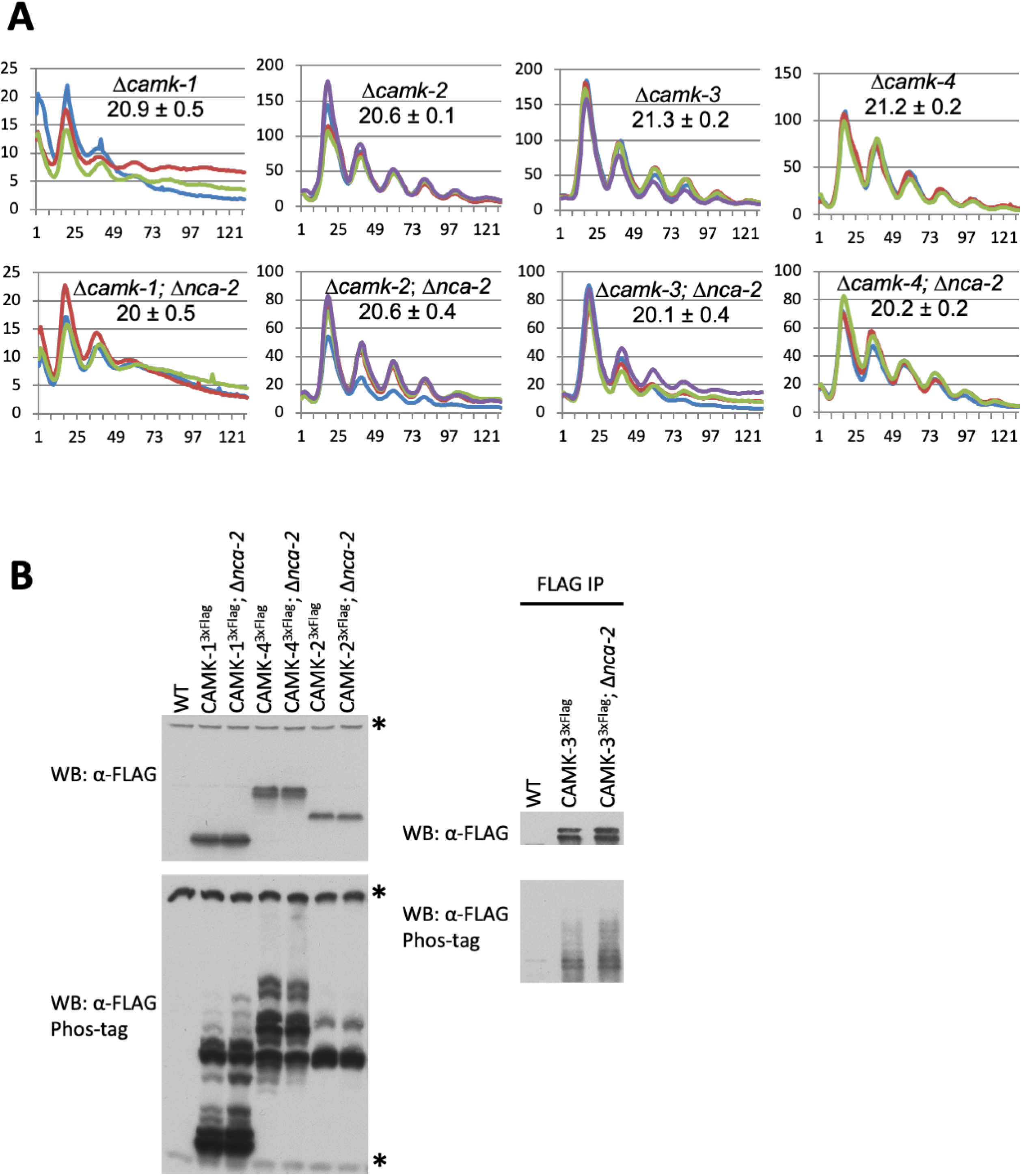
Period shortening of Δ*nca-2* is rescued by deletion of *camk-2*. (**A**) Luciferase assays were performed with a *frq C box* promoter-driven *luciferase* gene at the *his-3* in individual knockouts of *camks-1*-*4* in the presence or absence of Δ*nca-2* as indicated. Periods are reported as described in Material and Methods and is reported ± SD (n=3). (**B**) Upper panel, total levels of CAMKs-1-4 that have a 3xFLAG tag respectively at their C-termini in WT or the Δ*nca-2* background were assayed by Western blotting with FLAG antibody; lower panel, phosphorylation profiles of CAMKs-1-4 were analyzed for the same sample set with 149:1 acrylamide to bis-acrylamide SDS-PAGE gels containing Phos-tag. Asterisks indicate non-specific bands. For CAMKs-1, -2, and -4, total lysates were applied while CAMK-3 was first pulled down by FLAG antibody-conjugated resins and subsequently assayed in WB due to an overlap between CAMK-3 phospho-isoforms and non-specific bands in the Phos-tag gel.

### Characterization of *nca-2*

In the *Neurospora* genome, transcription of ∼40% coding genes are circadianly controlled directly or indirectly by the WCC-FFC oscillator (69, 70). We used transcriptional and translational fusions with the *luciferase* gene to see whether *nca-2* is a *ccg*: (1) the *nca-2* promoter was fused to the *luciferase* gene and transformed to the *csr* locus for real-time analysis of *nca-2* transcription, showing that transcription driven by the *nca-2* promoter is clearly rhythmic (Figure 5A); (2) after fusing the *nca-2* coding sequence with the *luciferase* open reading frame (ORF), tracking the bioluminescent signal of NCA-2-LUC protein reveals that the NCA-2-LUC signal also oscillates in a typical circadian manner (Figure 5B). These data indicate that calcium signaling in the cell might be regulated by the circadian clock through rhythmically transcribing and translating a calcium pump gene, *nca-2*. These data place NCA-2 in the larger cellular circadian system: *nca-2* and NCA-2 expression are clock-regulated, and NCA-2 activity, or lack thereof, impacts circadian period length. To identify potential DNA elements conferring circadian transcription of *nca-2*, we searched rhythmic motifs derived from (69). These were identified as sequences that were over-represented among rhythmically expressed genes. Interestingly, the first three of the four types of motifs identified in the paper (69) are found in the *nca-2* promoter (1.7 kb upstream of “ATG”) (data not shown). However, we do not know what TFs bind to these motifs; they do not appear in available databases, including the extensive catalogue of inferred sequence preferences of DNA-binding proteins (Cis-BP) (http://cisbp.ccbr.utoronto.ca) (71) that covers >1000 TFs from 131 species including *Neurospora*. Although there were weak matches to the motifs, none of the matches were from *Neurospora* (data not shown).

**Figure 5.**
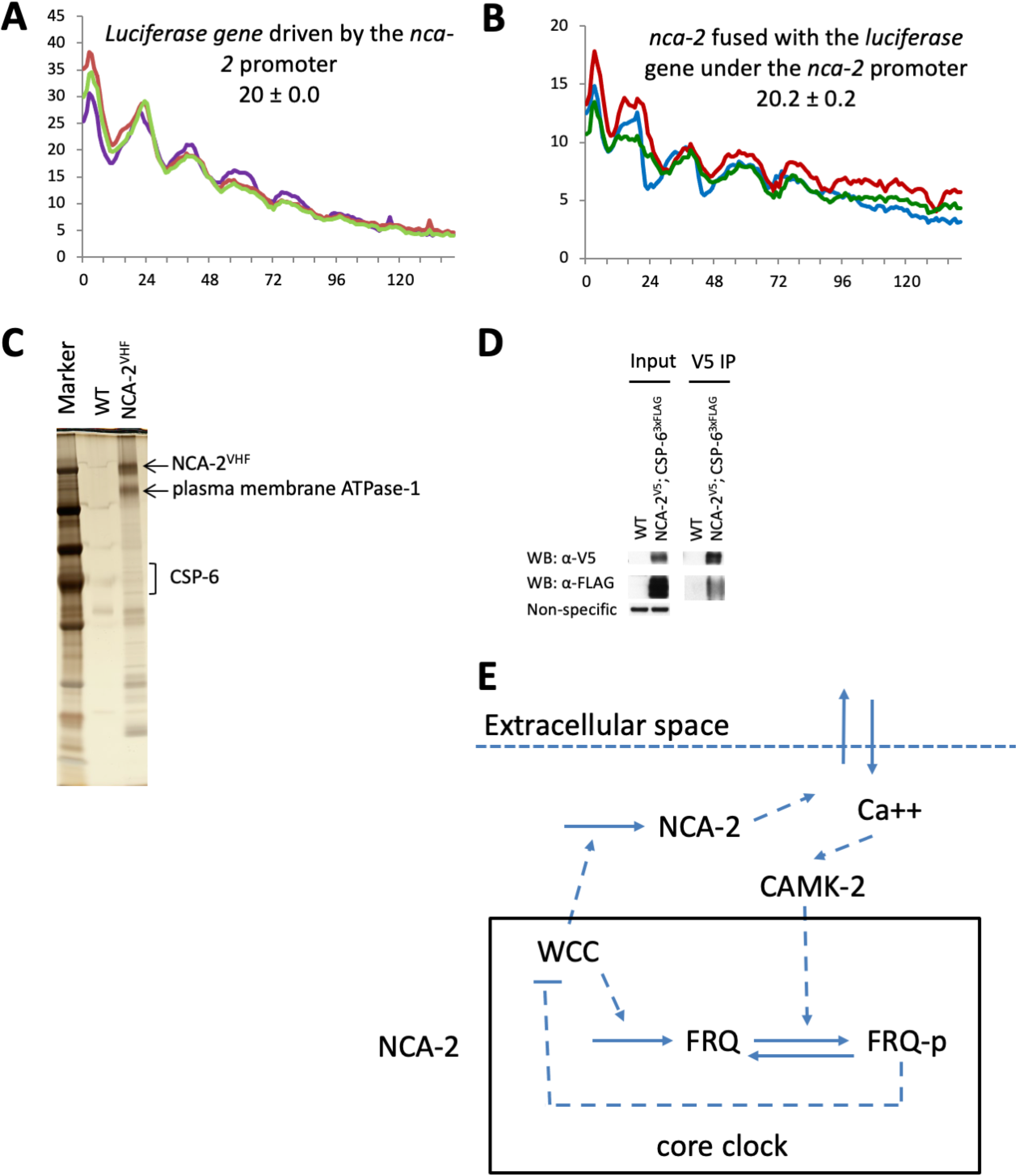
*nca-2* is a *ccg* and modulates both input to and output from the core clock. (**A**) The *nca-2* promoter fused to the *luciferase* gene was transformed to the *csr* locus and luciferase signals were followed at 25 °C in the dark. Period was determined as described in Material and Methods and is reported ± SD (n=3). (**B**) The *nca-2* open reading frame was fused to the 5’ end of the firefly *luciferase* gene and the same assay as in (**A**) was performed to trace the luciferase signal. (**C**) A representative silver-stained gel showing NCA-2^VHF^ and its interactome purified from a culture grown in the light. NCA-2^VHF^ and interactors were affinity-purified, trichloroacetic acid (TCA)-precipitated, and analyzed by mass spectrometry. (**D**) NCA-2 is tagged with a V5 tag and one of its interactors, CSP-6, was tagged with a 3xFLAG tag; Co-immunoprecipitation was performed using V5 resin and Western blotting was done with V5 and FLAG antibodies respectively. (**E**) Working model for the roles of intracellular calcium and of *nca-2* in the circadian system, In Δ*nca-2*, increased calcium over-activates CAMKs, which induces FRQ over-phosphorylations and thereby causes a faster running clock; the circadian clock regulates expression of *nca-2* and *camk*s.

Consistent with its role as a calcium exporter, NCA-2 is predicted to contain two calcium ATPase domains and a haloacid dehalogenase (HAD) domain (Figure S5A). To understand the role of NCA-2 at a mechanistic level, we mapped the NCA-2 interactome by affinity purification. C-terminal V5-10xHis-3xFLAG (VHF)-tagged NCA-2 was affinity-purified under a non-denaturing condition (Figure 5C), and its interacting proteins identified by mass spectrometry. Among NCA-2’s interactors identified (Table S1) was the phosphatase CSP-6 whose interaction with NCA-2 was confirmed by immuno-precipitation (Figure 5D). CSP-6 has been shown to control circadian output and WCC phosphorylations independent of the circadian feedback loop (72), suggesting that NCA-2 might have other roles relevant to CSP-6. Both Δ*csp-6* and the double mutant Δ*csp-6*; Δ*nca-2* display an arrhythmic overt clock on race tubes (Figure S5B), indicating that Δ*nca-2* is unable to rescue the output defect in Δ*csp-6*. Interestingly however, while growing more slowly than Δ*csp-6*, the double mutant Δ*csp-6*; Δ*nca-2* shows a period similar to Δ*nca-2* by the luciferase assay (Figure S5C), suggesting that *nca-2* does not act through *csp-6* in controlling the pace of the core oscillator. Altogether, these data demonstrate that *nca-2* is a *ccg* and suggest that cellular calcium signaling might be regulated by the circadian clock via rhythmic expression of *nca-2*.

### Downregulation of *calcineurin* does not influence the circadian period

In a wide variety of eukaryotes, a prolonged increase in intracellular Ca^2+^ activates a calcium and calmodulin dependent serine/threonine protein phosphatase, calcineurin, which mediates dephosphorylation of transcription factors, such as NFAT to regulate gene expression (73–86). *calcineurin* (*ncu03833*) is an essential gene in *Candida albicans* and *Neurospora* (87, 88), so to determine whether *calcineurin* influences the circadian clock, we downregulated its expression by replacing its native promoter with the *qa-2* promoter, an inducible promoter activated by quinic acid (QA). In the absence of QA, WC-1 is undetectable and FRQ is barely seen in *qa-2-*driven *calcineurin* (Figure S6A), consistent with a short period/arrhythmic clock observed in the strain (Figure S6B). To better examine this, we assayed rhythmicity at extremely low levels of inducer just sufficient for rhythmicity (10^−8^ M QA) or at high levels at or above WT expression (10^−2^ M QA). We found that period length was not proportional to the level of *calcineurin* expression at levels supporting any rhythmicity, and that even at vanishingly low *calcineurin* expression the core oscillator displays a period similar to that in WT, suggesting that the level of *calcineurin* does not determine the pace of the clock. This said, the severe reduction in WC-1 levels in the *qa-2*-driven *calcineurin* strain cultured without QA would be consistent with at least an indirect role for *calcineurin* in controlling WC-1 expression.

## Discussion

In this study, we have identified *nca-2* **as** a calcium pump involved in regulating circadian period length through CAMK-mediated FRQ phosphorylations. These data confirm that calcium signaling, a crucial regulatory pathway in mediating cellular and biochemical processes, must be well controlled for normal circadian period length determination. Most significantly, calcium signaling is now placed as an ancillary feedback loop within the larger circadian oscillatory system: The clock controls the expression of NCA-2 and thereby intracellular calcium levels and intracellular calcium, in turn, modulates the period length of the clock. In this regard the larger *Neurospora* circadian system is regulated by calcium in a manner reminiscent of that seen in the mammalian brain (e.g. (89)). As prolonged activation of signaling pathways is wasteful and harmful to the cell, the elevated cytosolic calcium in Δ*nca-2* over-activates CAMKs, leading to FRQ hyperphosphorylation and thereby causing a period defect (Figure 5E). The involvement of intracellular Ca^2+^ in the circadian system is further nuanced by the finding that expression of some *camk* genes is clock-controlled (Figure S3 and (69, 70)), so both the activator and effectors of calcium-induced regulation are clock-modulated and clock-affecting. This emphasizes the pervasive nature of both circadian and calcium control of the biology of the cell (Figure 5E).

Among calcium trafficking genes, *nca-2* encodes the major Ca^2+^ exporter (34). *Neurospora* encodes three transporter *nca* genes as well as the vacuolar calcium importer gene *cax*, but interestingly, only disruption of *nca-2* leads to a significant period change (Figure 1A), suggesting that NCA-2 plays a major role in lowering cytosolic calcium. Consistent with this, the calcium level in Δ*nca-2* has been reported to rise ∼9.3 times while it remains normal in Δ*nca-1* or Δ*nca-3* (36). It is possible that NCA-2 has higher affinity for Ca^2+^, is more abundant on the plasma membrane, or more efficient in transporting calcium than the other two NCAs.

Temporal FRQ phosphorylation, the core pacemaking mechanism in the circadian feedback loop, is mediated by multiple kinases including at least CKI, CKII, and CAMK-1 (9, 16, 18). Deletion of *camk-2* gene prevents the high intracellular Ca^+2^ from shortening the circadian period, indicating the dominant role in mediating the effect of calcium on the clock and making it a likely addition to the CAMKs active on the clock. Periods of several *frq* phosphorylation mutants, *frq*^*S72A, S73A, S76A*^, *frq*^*S538A, S540A*^ and *frq*^*S632A, S634A*^ (Figure 3), were not significantly altered in the background of Δ*nca-2*, and the domain where FRQ S72, S73, and S76 are located bears CAMK motifs (9), consistent with calcium-activated CAMK acting through these residues. Interestingly, although it is CAMK-1 that has been shown to directly phosphorylate FRQ *in vitro* (18) its loss here did not abrogate the effects of loss of NCA-2. It may be that the phosphosites targeted by different CAMKs on FRQ are distinct and have different effects on rhythmicity. Freshly germinated Δ*camk-1* displays a developmental defect (18, 37), whereas knockouts of the other three *camks* individually grow as robustly as WT (Figure S4A); however, the growth defect of Δ*camk-1* strains appears to rapidly revert back to normal after a few rounds of inoculation of Δ*camk-1* on new slants (18), suggesting that other CAMKs might be able to gradually compensate for the loss of *camk-1* over time.

WCC can be phosphorylated at over 90 sites and a small group of these is required for the closure of the circadian feedback loop (10). Interestingly, in Δ*nca-2*, WC-1 is hyperphosphorylated and more abundant than in WT despite a reduced *wc-1* RNA level (Figure 1A); this finding is consistent with a “black widow” model in which site-specific phosphorylation of transcription activators makes them inactive in driving transcription but more stable (90). However, lacking key phosphoresidues determining the feedback loop closure, *wc-1* mutants, such as *wc-1*^*S971A, S988A, S990A, S992A, S994A, S995A*^; *wc-2*^*S394A, S428A, S429A, S433A, S435A*^, show rhythms with short circadian period lengths due to elevated activity of WCC (10), whereas Δ*nca-2* strains, bearing hyperphosphorylated and more stable WC-1, also display a short period (Figure 1 A and 1B; Figure 2C and 2E). One possible explanation is that the hyper-phosphorylation of WC-1 in Δ*nca-2* occurs at residues regulating the circadian amplitude/output instead of those required for the feedback loop closure while the period shortening effect in Δ*nca-2* is caused by enhanced FRQ phosphorylation. WCC phosphoresidues can be briefly classified into two categories: the ones involved in the feedback loop closure and the other ones regulating the robustness of *frq* transcription (amplitude reflecting peak to trough in circadian cycles) (10). Key phosphomutants of *wcc* showed an additive effect with Δ*nca-2* on period length, suggesting that NCA-2 is not directly involved in the regulation of sites participating in the feedback loop closure but instead regulates WCC phosphoresidues relevant to the circadian amplitude.

## Material and methods

### Strains and culture conditions

328-4 (*ras-1*^*bd*^ *A*) was used as a wild-type strain in the race tube analyses and, 661-4a (*ras-1*^*bd*^ A), which bears the *frq C-box* fused to codon-optimized *luciferase* gene at the *his-3* locus, served as the wild type in luciferase assays. *Neurospora* transformation was performed as previously reported (91, 92). Medium in the race tube analyses contains 1 x Vogel’s salts, 0.17% arginine, 1.5% agar, 50 ng/mL biotin, and 0.1% glucose and liquid culture medium (LCM) contains 1 x Vogel’s, 0.5% arginine, 50 ng/mL biotin, and 2% glucose. Unless otherwise specified, race tubes were cultured in constant light for 16–24 hrs at 25 °C to synchronize strains and then transferred to the dark at 25 °C. The Vogeloid (10X) used to make the Ca^2+^-free medium in Figure 1D contains 100 mM NH_4_Cl, 20 mM MgCl_2_.6H_2_O, 100 mM KCl, 20 mM methionine, 50 ng/mL biotin, and 0.1% glucose (36).

### Bioluminescence assays

Luciferase assays were conducted as previously described (10). Briefly, strains with the *frq C-box luciferase* transcriptional reporter at the *his-3* locus were grown in 96 well plates bearing 0.1% glucose race tube medium having luciferin in constant light overnight (16-24 hrs) at 25 °C and then transferred to the dark at 25 °C to start circadian cycles. Bioluminescent signals were tracked by a CCD-camera every hour for 5 or more days. Luciferase data were extracted using the NIH ImageJ software with a custom macro, and circadian period lengths were manually determined.

### Protein lysate and Western blot (WB)

For WB, 15 mgs of whole-cell protein lysate were loaded per lane on a 3-8% tris-acetate or 6.5% tris-glycine (bearing Phos-tag) SDS gel (92). Custom-raised antibodies against WC-1, WC-2, FRQ, and FRH have been described previously (93–95). V5 antibody (Thermo Pierce) and FLAG antibody (M2, Sigma-Aldrich) were diluted 1:5000 for use as the primary antibody. To analyze the phosphorylation profiles of CAMKs, 20 μM Phos-tag chemical (ApexBio) was added to the homemade 6.5% Tris-glycine SDS-PAGE gel bearing a ratio of 149:1 acrylamide/bisacrylamide (10).

### Immunoprecipitation (IP)

IP was performed as previously described (91, 92). Briefly, 2 mgs of total protein were incubated with 20 μL of V5 agarose (Sigma-Aldrich) rotating at 4 °C for 2 hr. The agarose beads were washed with 1 mL of protein extraction buffer (50 mM HEPES [pH 7.4], 137 mM NaCl, 10% glycerol, 0.4% NP-40) twice and eluted with 50 μL of 5 x SDS sample buffer at 99 °C for 5 min.

### Other techniques

RNA extraction, reverse transcription (RT), and quantitative PCR were conducted as previous reports (72, 91). VHF-tagged NCA-2 was purified with the same method applied for isolation of C-terminal VHF-tagged WC-1 and mass spectrometry analyses were performed as previously described (72, 91). Data acquisition and analysis of luciferase runs were carried out as previously described (10).

## Acknowledgements

We thank the Fungal Genetics Stock Center for providing *Neurospora* strains. This work was supported by grants from the NIH to J.C.D. (R35GM118021), J.J.L.(R35GM118022), and S.A.G. (R01GM122846).

## Legends

**Figure S1** Stability of WC-1 and FRQ in WT and Δ*nca-2*. (**A**) Left, cycloheximide (CHX) was added to the *Neurospora* culture growing in the light at a final concentration of 40 μg/mL and tissues were harvested at indicated time points and assayed by Western blotting with WC-1- and FRQ-specific antibodies respectively; right, densitometric analyses of the blots on the left showing WC-1 and FRQ levels in WT and Δ*nca-2* treated with 40 μg/mL CHX for 0, 4, 8, or 12 hrs as indicated. (**B**) Left, FRQ stability in WT and Δ*nca-2* is measured using CHX-treated cultures sampled every 2 hrs over 14 hrs; right, densitometric analysis of FRQ on the left panel.

**Figure S2** Δ*nca-2* shortens period length in a long period *frq* allele and in *wcc* phospho-mutants (**A**) The activity of the *frq* promoter is measured by the *C-box-luc* bioluminescence in the background of *frq*^7^, *wc-1*^*S971A, S988A, S990A, S992A, S994A, S995A*^, or *wc-2*^*S433A*^ in the presence or absence of *nca-2* as indicated in the figure. (**B**) Phosphorylations of WC-1 S971 and S990 in the presence or absence of *nca-2* were assayed using a 6.5% SDS-PAGE gel bearing Phos-tag (for details, see Materials and Methods). Phosphorylation of WC-1 residues S971, S988, S990, S992, S994, and S995 is promoted by FRQ and essential for the closure of the circadian feedback loop (10).

**Figure S3** Expression of *camks* in WT and Δ*nca-2*. mRNA levels of *camks-1-4* in WT and Δ*nca-2* were assayed by RT-qPCR with samples grown under light or dark conditions as indicated. Expression of *camks* was normalized to that of *rac-1*.

**Figure S4** Individual or combinational deletion of *camk* genes does not dramatically impact the circadian period by the luciferase assay.(**A**) Gene knockouts of individual *camks-1-4* were assayed for circadian phenotypes by race tube. Δ*camk-1* cultures in the top and middle tubes are true knockouts while the one at the bottom is a typical revertant as reported by (18) (**B**) Combinations of Δ*camk* mutants were tested by the luciferase assay.

**Figure S5** NCA-2 domain analyses and its relationship with *csp-6* on the clock. (**A**) Predicted domains in NCA-2 analyzed by the on-line tool SMART (http://smart.embl-heidelberg.de/smart/set_mode.cgi?NORMAL=1) Race tube (**B**) and luciferase (**C**) analyses of Δ*nca-2*, Δ*csp-6*, and Δ*csp-6*; Δ*nca-2*.

**Figure S6** Down-regulation of *calcineurin subunit B* has little influence on circadian period length across expression levels compatible with rhythmicity. The endogenous promoter of *calcineurin subunit B* was replaced by the *qa-2* promoter. (**A**) FRQ, WC-1, and WC-2 proteins were determined by Western blotting with specific antibodies and (**B**) *frq* transcription was tracked by *frq C box-luc* at the *his-3* locus in the absence or presence of quinic acid (QA) at the concentration indicated.

## References

1. Matsu-Ura T, Moore SR, Hong CI. 2018. WNT Takes Two to Tango: Molecular Links between the Circadian Clock and the Cell Cycle in Adult Stem Cells. J Biol Rhythms 33:5–14.

2. Haspel JA, Anafi R, Brown MK, Cermakian N, Depner C, Desplats P, Gelman AE, Haack M, Jelic S, Kim BS, Laposky AD, Lee YC, Mongodin E, Prather AA, Prendergast BJ, Reardon C, Shaw AC, Sengupta S, Szentirmai É, Thakkar M, Walker WE, Solt LA. 2020. Perfect timing: circadian rhythms, sleep, and immunity - an NIH workshop summary. JCI Insight 5.

3. Hevia MA, Canessa P, Larrondo LF. 2016. Circadian clocks and the regulation of virulence in fungi: Getting up to speed. Semin Cell Dev Biol 57:147–155.

4. Froehlich AC, Liu Y, Loros JJ, Dunlap JC. 2002. White Collar-1, a Circadian Blue Light Photoreceptor, Binding to the frequency Promoter. Science 297:815–819.

5. Froehlich AC, Loros JJ, Dunlap JC. 2003. Rhythmic binding of a WHITE COLLAR-containing complex to the frequency promoter is inhibited by FREQUENCY. Proceedings of the National Academy of Sciences 100:5914–5919.

6. He Q, Cheng P, Yang Y hong, Wang L, Gardner KH, Liu Y. 2002. White Collar-1, a DNA Binding Transcription Factor and a Light Sensor. Science 297:840–843.

7. Cheng P, He Q, He Q, Wang L, Liu Y. 2005. Regulation of the Neurospora circadian clock by an RNA helicase. Genes & Development 19:234–241.

8. Shi M, Collett M, Loros JJ, Dunlap JC. 2010. FRQ-Interacting RNA Helicase Mediates Negative and Positive Feedback in the Neurospora Circadian Clock. Genetics 184:351–361.

9. Baker CL, Kettenbach AN, Loros JJ, Gerber SA, Dunlap JC. 2009. Quantitative Proteomics Reveals a Dynamic Interactome and Phase-Specific Phosphorylation in the Neurospora Circadian Clock. Molecular Cell 34:354–363.

10. Wang B, Kettenbach AN, Zhou X, Loros JJ, Dunlap JC. 2019. The phospho-code determining circadian feedback loop closure and output in Neurospora. Molecular cell 74:771–784.

11. Lipton JO, Yuan ED, Boyle LM, Ebrahimi-Fakhari D, Kwiatkowski E, Nathan A, Güttler T, Davis F, Asara JM, Sahin M. 2015. The Circadian Protein BMAL1 Regulates Translation in Response to S6K1-Mediated Phosphorylation. Cell 161:1138–1151.

12. Luciano AK, Zhou W, Santana JM, Kyriakides C, Velazquez H, Sessa WC. 2018. CLOCK phosphorylation by AKT regulates its nuclear accumulation and circadian gene expression in peripheral tissues. J Biol Chem 293:9126–9136.

13. Narasimamurthy R, Hunt SR, Lu Y, Fustin J-M, Okamura H, Partch CL, Forger DB, Kim JK, Virshup DM. 2018. CK1δ /ε protein kinase primes the PER2 circadian phosphoswitch. Proc Natl Acad Sci USA 115:5986–5991.

14. Robles MS, Humphrey SJ, Mann M. 2017. Phosphorylation Is a Central Mechanism for Circadian Control of Metabolism and Physiology. Cell Metabolism 25:118–127.

15. Tang CT, Li S, Long C, Cha J, Huang G, Li L, Chen S, Liu Y. 2009. Setting the pace of the Neurospora circadian clock by multiple independent FRQ phosphorylation events. Proceedings of the National Academy of Sciences 106:10722–10727.

16. He Q, Cha J, He Q, Lee H-C, Yang Y, Liu Y. 2006. CKI and CKII mediate the FREQUENCY-dependent phosphorylation of the WHITE COLLAR complex to close the Neurospora circadian negative feedback loop. Genes & Development 20:2552–2565.

17. Huang G, Chen S, Li S, Cha J, Long C, Li L, He Q, Liu Y. 2007. Protein kinase A and casein kinases mediate sequential phosphorylation events in the circadian negative feedback loop. Genes & Development 21:3283–3295.

18. Yang Y, Cheng P, Zhi G, Liu Y. 2001. Identification of a Calcium/Calmodulin-dependent Protein Kinase That Phosphorylates the Neurospora Circadian Clock Protein FREQUENCY. J Biol Chem 276:41064–41072.

19. Schafmeier T, Haase A, Káldi K, Scholz J, Fuchs M, Brunner M. 2005. Transcriptional Feedback of Neurospora Circadian Clock Gene by Phosphorylation-Dependent Inactivation of Its Transcription Factor. Cell 122:235–246.

20. Schwerdtfeger C, Linden H. 2000. Localization and light-dependent phosphorylation of white collar 1 and 2, the two central components of blue light signaling in Neurospora crassa: Blue light signal transduction in N. crassa. European Journal of Biochemistry 267:414–422.

21. Duchen MR. 2012. Mitochondria, calcium-dependent neuronal death and neurodegenerative disease. Pflugers Arch - Eur J Physiol 464:111–121.

22. Berchtold MW, Brinkmeier H, Müntener M. 2000. Calcium Ion in Skeletal Muscle: Its Crucial Role for Muscle Function, Plasticity, and Disease. Physiological Reviews 80:1215–1265.

23. Wakai T, Vanderheyden V, Fissore RA. 2011. Ca2+ Signaling During Mammalian Fertilization: Requirements, Players, and Adaptations. Cold Spring Harbor Perspectives in Biology 3:a006767– a006767.

24. Lee MJ, Yaffe MB. 2016. Protein Regulation in Signal Transduction. Cold Spring Harb Perspect Biol 8:a005918.

25. Love J, Dodd AN, Webb AAR. 2004. Circadian and diurnal calcium oscillations encode photoperiodic information in Arabidopsis. Plant Cell 16:956–966.

26. Johnson CH, Knight MR, Kondo T, Masson P, Sedbrook J, Haley A, Trewavas A. 1995. Circadian oscillations of cytosolic and chloroplastic free calcium in plants. Science 269:1863– 1865.

27. MartíRuiz MC, Hubbard KE, Gardner MJ, Jung HJ, Aubry S, Hotta CT, Mohd-Noh NI, Robertson FC, Hearn TJ, Tsai Y-C, Dodd AN, Hannah M, Carré IA, Davies JM, Braam J, Webb AAR. 2018. Circadian oscillations of cytosolic free calcium regulate the Arabidopsis circadian clock. Nat Plants 4:690–698.

28. Imaizumi T, Schroeder JI, Kay SA. 2007. In SYNC: The ins and outs of circadian oscillations in calcium. Sci STKE 2007:pe32.

29. Techel D, Gebauer G, Kohler W, Braumann T, Jastorff B, Rensing L. 1990. On the role of Ca2(+)-calmodulin-dependent and cAMP-dependent protein phosphorylation in the circadian rhythm of Neurospora crassa. J Comp Physiol B 159:695–706.

30. Nakashima H. 1984. Calcium Inhibits Phase Shifting of the Circadian Conidiation Rhythm of Neurospora crassa by the Calcium Ionophore A23187. Plant Physiol 74:268–271.

31. Nakashima H. 1986. Phase shifting of the circadian conidiation rhythm in Neurospora crassa by calmodulin antagonists. J Biol Rhythms 1:163–169.

32. Sadakane Y, Nakashima H. 1996. Light-induced phase shifting of the circadian conidiation rhythm is inhibited by calmodulin antagonists in Neurospora crassa. J Biol Rhythms 11:234–240.

33. Suzuki S, Katagiri S, Nakashima H. 1996. Mutants with altered sensitivity to a calmodulin antagonist affect the circadian clock in Neurospora crassa. Genetics 143:1175–1180.

34. Bowman BJ, Draskovic M, Freitag M, Bowman EJ. 2009. Structure and Distribution of Organelles and Cellular Location of Calcium Transporters in Neurospora crassa. Eukaryotic Cell 8:1845–1855.

35. Bowman BJ, Abreu S, Johl JK, Bowman EJ. 2012. The pmr Gene, Encoding a Ca2+-ATPase, Is Required for Calcium and Manganese Homeostasis and Normal Development of Hyphae and Conidia in Neurospora crassa. Eukaryot Cell 11:1362–1370.

36. Bowman BJ, Abreu S, Margolles-Clark E, Draskovic M, Bowman EJ. 2011. Role of Four Calcium Transport Proteins, Encoded by nca-1, nca-2, nca-3, and cax, in Maintaining Intracellular Calcium Levels in Neurospora crassa. Eukaryotic Cell 10:654–661.

37. Kumar R, Tamuli R. 2014. Calcium/calmodulin-dependent kinases are involved in growth, thermotolerance, oxidative stress survival, and fertility in Neurospora crassa. Arch Microbiol 196:295–305.

38. Laxmi V, Tamuli R. 2017. The calmodulin gene in Neurospora crassa is required for normal vegetative growth, ultraviolet survival, and sexual development. Arch Microbiol 199:531–542.

39. Deka R, Tamuli R. 2013. Neurospora crassa ncs-1, mid-1 and nca-2 double-mutant phenotypes suggest diverse interaction among three Ca(2+)-regulating gene products. J Genet 92:559–563.

40. Barman A, Tamuli R. 2017. The pleiotropic vegetative and sexual development phenotypes of Neurospora crassa arise from double mutants of the calcium signaling genes plc-1, splA2, and cpe-1. Curr Genet 63:861–875.

41. Cui J, Kaandorp JA, Sloot PMA, Lloyd CM, Filatov MV. 2009. Calcium homeostasis and signaling in yeast cells and cardiac myocytes. FEMS Yeast Res 9:1137–1147.

42. Liu S, Hou Y, Liu W, Lu C, Wang W, Sun S. 2015. Components of the Calcium-Calcineurin Signaling Pathway in Fungal Cells and Their Potential as Antifungal Targets. Eukaryot Cell 14:324–334.

43. Cui J, Kaandorp JA, Ositelu OO, Beaudry V, Knight A, Nanfack YF, Cunningham KW. 2009. Simulating calcium influx and free calcium concentrations in yeast. Cell Calcium 45:123–132.

44. Endo M. 2009. Calcium-induced calcium release in skeletal muscle. Physiol Rev 89:1153– 1176.

45. Jones HC, Keep RF. 1987. The control of potassium concentration in the cerebrospinal fluid and brain interstitial fluid of developing rats. J Physiol 383:441–453.

46. Berg JM, Tymoczko JL, Stryer L. 2002. Calcium Ion Is a Ubiquitous Cytosolic Messenger. Biochemistry 5th edition.

47. Alzheimer C. 2013. Na Channels and Ca2+ Channels of the Cell Membrane as Targets of Neuroprotective SubstancesMadame Curie Bioscience Database [Internet]. Landes Bioscience.

48. Bowman BJ, Dschida WJ, Bowman EJ. 1992. Vacuolar ATPase of Neurospora crassa: electron microscopy, gene characterization and gene inactivation/mutation. J Exp Biol 172:57–66.

49. Bowman EJ, Bowman BJ. 2000. Cellular role of the V-ATPase in Neurospora crassa: analysis of mutants resistant to concanamycin or lacking the catalytic subunit A. J Exp Biol 203:97–106.

50. Margolles-Clark E, Tenney K, Bowman EJ, Bowman BJ. 1999. The structure of the vacuolar ATPase in Neurospora crassa. J Bioenerg Biomembr 31:29–37.

51. Bowman BJ, Vázquez-Laslop N, Bowman EJ. 1992. The vacuolar ATPase of Neurospora crassa. J Bioenerg Biomembr 24:361–370.

52. Yáñez M, Gil-Longo J, Campos-Toimil M. 2012. Calcium binding proteins. Adv Exp Med Biol 740:461–482.

53. Zamponi GW, Currie KPM. 2013. Regulation of CaV2 calcium channels by G protein coupled receptors. Biochimica et Biophysica Acta (BBA) - Biomembranes 1828:1629–1643.

54. Predescu Crețoiu, Creț oiu, Pavelescu, Suciu, Radu, Voinea. 2019. G Protein-Coupled Receptors (GPCRs)-Mediated Calcium Signaling in Ovarian Cancer: Focus on GPCRs activated by Neurotransmitters and Inflammation-Associated Molecules. IJMS 20:5568.

55. Swulius MT, Waxham MN. 2008. Ca2+/Calmodulin-dependent Protein Kinases. Cell Mol Life Sci 65:2637–2657.

56. Lee K-T, So Y-S, Yang D-H, Jung K-W, Choi J, Lee D-G, Kwon H, Jang J, Wang LL, Cha S, Meyers GL, Jeong E, Jin J-H, Lee Y, Hong J, Bang S, Ji J-H, Park G, Byun H-J, Park SW, Park Y-M, Adedoyin G, Kim T, Averette AF, Choi J-S, Heitman J, Cheong E, Lee Y-H, Bahn Y-S. 2016. Systematic functional analysis of kinases in the fungal pathogen Cryptococcus neoformans. 1. Nature Communications 7:12766.

57. Gustin MC, Albertyn J, Alexander M, Davenport K. 1998. MAP Kinase Pathways in the Yeast Saccharomyces cerevisiae. Microbiol Mol Biol Rev 62:1264–1300.

58. Joseph JD, Means AR. 2002. Calcium Binding Is Required for Calmodulin Function in Aspergillus nidulans. Eukaryot Cell 1:119–125.

59. Lévy J, Bres C, Geurts R, Chalhoub B, Kulikova O, Duc G, Journet E-P, Ané J-M, Lauber E, Bisseling T, Dénarié J, Rosenberg C, Debellé F. 2004. A Putative Ca2+ and Calmodulin-Dependent Protein Kinase Required for Bacterial and Fungal Symbioses. Science 303:1361– 1364.

60. Soderling TR. 1999. The Ca2+–calmodulin-dependent protein kinase cascade. Trends in Biochemical Sciences 24:232–236.

61. Ohya Y, Kawasaki H, Suzuki K, Londesborough J, Anraku Y. 1991. Two yeast genes encoding calmodulin-dependent protein kinases. Isolation, sequencing and bacterial expressions of CMK1 and CMK2. J Biol Chem 266:12784–12794.

62. Tsai P-J, Tu J, Chen T-H. 2002. Cloning of a Ca(2+)/calmodulin-dependent protein kinase gene from the filamentous fungus Arthrobotrys dactyloides. FEMS Microbiol Lett 212:7–13.

63. Takeda N, Maekawa T, Hayashi M. 2012. Nuclear-localized and deregulated calcium- and calmodulin-dependent protein kinase activates rhizobial and mycorrhizal responses in Lotus japonicus. Plant Cell 24:810–822.

64. Chen J, Gutjahr C, Bleckmann A, Dresselhaus T. 2015. Calcium Signaling during Reproduction and Biotrophic Fungal Interactions in Plants. Molecular Plant 8:595–611.

65. Wang J-P, Munyampundu J-P, Xu Y-P, Cai X-Z. 2015. Phylogeny of Plant Calcium and Calmodulin-Dependent Protein Kinases (CCaMKs) and Functional Analyses of Tomato CCaMK in Disease Resistance. Front Plant Sci 6:1075.

66. Impey S, Fong AL, Wang Y, Cardinaux J-R, Fass DM, Obrietan K, Wayman GA, Storm DR, Soderling TR, Goodman RH. 2002. Phosphorylation of CBP Mediates Transcriptional Activation by Neural Activity and CaM Kinase IV. Neuron 34:235–244.

67. Freitag SI, Wong J, Young PG. 2014. Genetic and physical interaction of Ssp1 CaMKK and Rad24 14-3-3 during low pH and osmotic stress in fission yeast. Open Biol 4.

68. Kinoshita E, Kinoshita-Kikuta E, Takiyama K, Koike T. 2006. Phosphate-binding tag, a new tool to visualize phosphorylated proteins. Mol Cell Proteomics 5:749–757.

69. Hurley JM, Dasgupta A, Emerson JM, Zhou X, Ringelberg CS, Knabe N, Lipzen AM, Lindquist EA, Daum CG, Barry KW, Grigoriev IV, Smith KM, Galagan JE, Bell-Pedersen D, Freitag M, Cheng C, Loros JJ, Dunlap JC. 2014. Analysis of clock-regulated genes in Neurospora reveals widespread posttranscriptional control of metabolic potential. Proc Natl Acad Sci USA 111:16995– 17002.

70. Hurley JM, Jankowski MS, De Los Santos H, Crowell AM, Fordyce SB, Zucker JD, Kumar N, Purvine SO, Robinson EW, Shukla A, Zink E, Cannon WR, Baker SE, Loros JJ, Dunlap JC. 2018. Circadian Proteomic Analysis Uncovers Mechanisms of Post-Transcriptional Regulation in Metabolic Pathways. Cell Syst 7:613-626.e5.

71. Weirauch MT, Yang A, Albu M, Cote AG, Montenegro-Montero A, Drewe P, Najafabadi HS, Lambert SA, Mann I, Cook K, Zheng H, Goity A, van Bakel H, Lozano J-C, Galli M, Lewsey MG, Huang E, Mukherjee T, Chen X, Reece-Hoyes JS, Govindarajan S, Shaulsky G, Walhout AJM, Bouget F-Y, Ratsch G, Larrondo LF, Ecker JR, Hughes TR. 2014. Determination and Inference of Eukaryotic Transcription Factor Sequence Specificity. Cell 158:1431–1443.

72. Zhou X, Wang B, Emerson JM, Ringelberg CS, Gerber SA, Loros JJ, Dunlap JC. 2018. A HAD family phosphatase CSP-6 regulates the circadian output pathway in Neurospora crassa. PLoS Genet 14:e1007192.

73. Park Y-J, Yoo S-A, Kim M, Kim W-U. 2020. The Role of Calcium–Calcineurin–NFAT Signaling Pathway in Health and Autoimmune Diseases. Front Immunol 11.

74. Juvvadi PR, Lee SC, Heitman J, Steinbach WJ. 2016. Calcineurin in fungal virulence and drug resistance: Prospects for harnessing targeted inhibition of calcineurin for an antifungal therapeutic approach. Virulence 8:186–197.

75. Park H-S, Lee SC, Cardenas ME, Heitman J. 2019. Calcium-calmodulin-calcineurin signaling: A globally conserved virulence cascade in eukaryotic microbial pathogens. Cell Host Microbe 26:453–462.

76. Thewes S. 2014. Calcineurin-Crz1 Signaling in Lower Eukaryotes. Eukaryot Cell 13:694–705.

77. Juvvadi PR, Lamoth F, Steinbach WJ. 2014. Calcineurin as a Multifunctional Regulator: Unraveling Novel Functions in Fungal Stress Responses, Hyphal Growth, Drug Resistance, and Pathogenesis. Fungal Biol Rev 28:56–69.

78. Stie J, Fox D. 2008. Calcineurin Regulation in Fungi and Beyond. Eukaryot Cell 7:177–186.

79. Vellanki S, Billmyre RB, Lorenzen A, Campbell M, Turner B, Huh EY, Heitman J, Lee SC. 2020. A Novel Resistance Pathway for Calcineurin Inhibitors in the Human-Pathogenic Mucorales Mucor circinelloides. mBio 11.

80. Loss O, Bertuzzi M, Yan Y, Fedorova N, McCann BL, Armstrong-James D, Espeso EA, Read ND, Nierman WC, Bignell EM. 2017. Mutual independence of alkaline- and calcium-mediated signalling in Aspergillus fumigatus refutes the existence of a conserved druggable signalling nexus. Mol Microbiol 106:861–875.

81. Bendickova K, Tidu F, Fric J. 2017. Calcineurin–NFAT signalling in myeloid leucocytes: new prospects and pitfalls in immunosuppressive therapy. EMBO Mol Med 9:990–999.

82. Steinbach WJ, Cramer RA, Perfect BZ, Asfaw YG, Sauer TC, Najvar LK, Kirkpatrick WR, Patterson TF, Benjamin DK, Heitman J, Perfect JR. 2006. Calcineurin Controls Growth, Morphology, and Pathogenicity in Aspergillus fumigatus. Eukaryot Cell 5:1091–1103.

83. Kozubowski L, Aboobakar EF, Cardenas ME, Heitman J. 2011. Calcineurin Colocalizes with P-Bodies and Stress Granules during Thermal Stress in Cryptococcus neoformans ▿. Eukaryot Cell 10:1396–1402.

84. Hill JA, Ammar R, Torti D, Nislow C, Cowen LE. 2013. Genetic and Genomic Architecture of the Evolution of Resistance to Antifungal Drug Combinations. PLoS Genet 9.

85. Aramburu J, Heitman J, Crabtree GR. 2004. Calcineurin: a central controller of signalling in eukaryotes. EMBO Rep 5:343–348.

86. Cruz MC, Fox DS, Heitman J. 2001. Calcineurin is required for hyphal elongation during mating and haploid fruiting in Cryptococcus neoformans. EMBO J 20:1020–1032.

87. Blankenship JR, Wormley FL, Boyce MK, Schell WA, Filler SG, Perfect JR, Heitman J. 2003. Calcineurin Is Essential for Candida albicans Survival in Serum and Virulence. Eukaryot Cell 2:422–430.

88. Tamuli R, Deka R, Borkovich KA. 2016. Calcineurin Subunits A and B Interact to Regulate Growth and Asexual and Sexual Development in Neurospora crassa. PLoS One 11.

89. Hastings MH, Brancaccio M, Maywood ES. 2014. Circadian pacemaking in cells and circuits of the suprachiasmatic nucleus. J Neuroendocrinol 26:2–10.

90. Tansey WP. 2001. Transcriptional activation: risky business. Genes & Development 15:1045–1050.

91. Wang B, Kettenbach AN, Gerber SA, Loros JJ, Dunlap JC. 2014. Neurospora WC-1 Recruits SWI/SNF to Remodel frequency and Initiate a Circadian Cycle. PLOS Genetics 10:e1004599.

92. Wang B, Zhou X, Loros JJ, Dunlap JC. 2016. Alternative Use of DNA Binding Domains by the Neurospora White Collar Complex Dictates Circadian Regulation and Light Responses. Mol Cell Biol 36:781–793.

93. Lee K, Loros JJ, Dunlap JC. 2000. Interconnected feedback loops in the Neurospora circadian system. Science 289:107–110.

94. Garceau NY, Liu Y, Loros JJ, Dunlap JC. Alternative Initiation of Translation and Time-Specific Phosphorylation Yield Multiple Forms of the Essential Clock Protein FREQUENCY 8.

95. Denault DL, Loros JJ, Dunlap JC. 2001. WC-2 mediates WC-1-FRQ interaction within the PAS protein-linked circadian feedback loop of Neurospora. EMBO J 20:109–117.

